# Primary care use of laboratory tests in Northern Ireland’s Western Health and Social Care Trust: a cross-sectional study

**DOI:** 10.1101/573949

**Authors:** Magda Bucholc, Maurice J O’Kane, Ciaran Mullan, Siobhan Ashe, Liam Maguire

## Abstract

**Objectives:** To describe the laboratory test ordering patterns by general practitioners (GPs) in Northern Ireland Western Health and Social Care Trust (WHSCT) and establish demographic and socio-economic associations with test requesting.

**Design:** Cross-sectional study.

**Setting:** Western Health and Social Care Trust, Northern Ireland

**Participants:** 55 WHSCT general practices requesting laboratory tests in the period from 1 April 2011 to 31 March 2016

**Outcomes:** To identify the temporal patterns of laboratory test ordering behaviour for 8 commonly requested clinical biochemistry tests/test groups in WHSCT. To analyse the extent of variations in laboratory test requests by GPs and to determine whether these variations can be accounted for by clinical outcomes or geographical, demographic, and socioeconomic characteristics.

**Results:** We identified substantial changes in the median number of request rates over five consecutive years of the study period as well as a large variation of adjusted test request rates for individual tests (lowest for electrolyte profiles, liver profiles, and HbA_1c_ and highest for immunoglobulins). No statistically significant relationship between ordering activity and either demographic (age and gender) and socioeconomic factors (deprivation) or Quality and Outcome Framework (QOF) scores was observed. We found that practice setting accounted for some of the between-practice variation in test requesting. Rural practices were characterized by both higher between practice variability and median number of order tests than urban practices at all time points.

**Conclusions:** A large between-practice variation in GP laboratory test requesting appears unrelated to demographic and socioeconomic indicators of the practices or crude clinical outcome indicators, most likely reflects differences in the clinical practice of individuals, potentially amenable to change through clinical interventions.

**Strengths and limitations of this study:**

- The study provides a comprehensive analysis of temporal changes in laboratory test utilization patterns and establishes the extent of variability in test requesting activity across general practices in Northern Ireland’s Western Health and Social Care Trust.
- The substantial variation in test ordering, not related to demographic and socioeconomic characteristics of practices, practice location or clinical outcome indicators, may reflect inappropriate laboratory test utilization and hence, suggest a potential for more efficient demand management of laboratory services.
- Given a cohort of general practices within one catchment area, our results provide evidence of differences in behaviour of individual GPs when managing patients with similar clinical symptoms.
- Failure to collect and cross-tabulate data on characteristics of general practitioners (GPs), such as GP’s age, years of experience, medical training was a study limitation and a missed opportunity in assessing the influence of practitioner factors on the variation in test ordering behaviour.

## Introduction

Despite the important role of laboratory testing in the diagnosis and monitoring of disease, there is concern about the increasing number of requested tests and in particular, large differences in laboratory utilization between clinical teams.^1^ In the UK, laboratory test orders grew by approximately 5% per year in recent years and inappropriate test requests are considered to be an important cause for this increase^2,3^. Although pathology expenditures account for only 5–6% of the UK total health budget, they are viewed as a potential source of savings, most likely because the costs can be easily identified and measured.^4^ According to the Department of Health, the rationalization of pathology services, including demand management of laboratory tests and elimination of unnecessary requesting, could produce savings of at least £500 million.^5^

Unnecessary testing is not only wasteful of resources but impacts on patients directly through the requirement for venepuncture and the follow up of minor (and possibly insignificant) abnormalities which may cause patient anxiety. On the other hand, inappropriate under requesting may cause harm through failure to diagnose or manage disease optimally. Several studies suggested that unnecessary and inappropriate utilization of laboratory services is closely linked to inter-practitioner variability in test requesting.^6-8^ Despite the increased availability of clinical management guidelines promoting harmonization of the use of laboratory tests, there is still substantial variation in test utilization among general practitioners.^7,9^ These differences appear to be unrelated to demographic characteristics of patient populations, socio-economic status of GP practices, disease prevalence or clinical outcome indicators.^6,7,10,11^ Even if some of these variables have been shown to have an effect on test ordering patterns, the variation in requesting rates is so large that it can only be explained by differences in attitudes towards the use of laboratory tests of individual practitioners.^10^ Accordingly, factors such as confidence in clinical judgement, clinical experience, an attitude about clinical practice guidelines, a lack of knowledge regarding the correct use of tests and fear of litigation have been identified as potential sources of practice variation.^12-15^

Since unwarranted variation can lead to suboptimal clinical outcomes,^16,17^ identification of factors contributing to differences in test requesting can provide useful information for optimising utilization of laboratory services. The aim of our study was to establish the extent of variability in test requesting and characterise temporal changes in test ordering patterns across general practices within the catchment area of the Northern Ireland (NI) Western Health and Social Care Trust (WHSCT) for a range of commonly requested clinical biochemistry tests/test groups. In addition, we investigated potential factors associated with inter-doctor variability in the use of laboratory tests including geographical, demographic, and socioeconomic factors as well as Quality and Outcome Framework (QOF) scores.

## Materials and Methods

### Study design and data sources

We conducted a cross-sectional study of laboratory test ordering activity across general practices in the WHSCT in the period from 1 April 2011 to 31 March 2016. The data on the use of laboratory tests were obtained from a HSC Business Object XI clinical information system. We investigated requesting rates for 8 clinical biochemistry tests/test groups including electrolyte profile, thyroid profile (FT4 and TSH), liver profile, lipid profile, urine albumin/creatinine (ACR) ratio, glycosylated haemoglobin (HbA_1c_), prostate-specific antigen (PSA), and immunoglobulins. The number of laboratory tests requested in each general practice was normalized by the number of registered patients and expressed as requests per 1000 patients, with the exception of H_b_A_1c_ and ACR for which ordering rates were expressed as tests per patient with diabetes and PSA rates, calculated per 1000 male patients. GP practice list size data (including the number of patients with diabetes) and patient demographic data was extracted from the Business Services Organisation (BSO) Family Practitioner Service Information and Registration Unit system. The rural-urban distribution of GP practices was based on the data from the Census Office of the Northern Ireland Statistics and Research Agency (NISRA).^18^ Socioeconomic characteristics of GP practices were determined using the NISRA Neighbourhood Information System (NINIS).^19^ QOF scores for individual practices were extracted from the website of Northern Ireland Department of Health.^20^

### Participants and setting

Data on laboratory tests requested from 55 general practices within the WHSCT were collected from the laboratory databases of Clinical Chemistry departments of the Altnagelvin Area Hospital, Tyrone County Hospital, and the Erne Hospital (subsequently the South West Acute Hospital). WHSCT provides health and social care services throughout the west of Northern Ireland, across the council areas of Derry City and Strabane District Council, Limavady in the Causeway Coast and Glens Borough Council, and Fermanagh and Omagh District Council.

### Inclusion criteria

We examined only laboratory test requests from 55 separate primary care medical practices within the catchment area of WHSCT that remained open throughout the study period i.e. during five consecutive years: Apr 2011 – Mar 2012, Apr 2012 – Mar 2013, Apr 2013 – Mar 2014, Apr 2014 – Mar 2015, and Apr 2015 – Mar 2016, were examined. To ensure the completeness and consistency of the data, test orders from WHSCT practices that closed or merged during the period of investigation were not taken into account.

### Variables and characteristics

Our analysis was limited to 8 frequently requested clinical biochemistry tests/test groups including electrolyte profile, thyroid profile, liver profile, lipid profile, ACR, HbA_1c_, PSA, and immunoglobulins. All considered tests/profiles (with the exception of immunoglobulins and ACR) were listed on a paper pathology request form and ordered by ticking a box adjacent to the test name; requests for ACR and immunoglobulins were made by writing the test name in an open text field.

Rates of test requests were analysed in the context of geographical and socioeconomic characteristics of practices, patient demographics (age and gender), and clinical outcome indicators. Note that we were not able to obtain a consolidated data on both gender and age of patients registered in individual GP practices.

The practice setting (rural v. urban area) was determined using the urban-rural classification of the Department for Environment, Food and Rural Affairs (DEFRA)^21^ while the size of settlements was obtained from the NISRA Census Office.^18^ A GP practice was defined as rural if its physical address was situated in a settlement of less than 10,000. Accordingly, we classified 24 practices as rural and 31 as urban. Furthermore, GP practices were categorised based on the Northern Ireland Multiple Deprivation Measure (MDM) identified for individual Super Output Areas (SOAs).^19^ The MDM comprises a weighted combination of 7 component measures (Income Deprivation; Employment Deprivation; Health Deprivation and Disability; Education, Skills and Training Deprivation; Proximity to Services; Living Environment; and Crime and Disorder) and ranges from 1 (most deprived SOAs) to 890 (least deprived SOAs). The Health Deprivation & Disability rank (one of the MDM domains)^19^ was analysed separately as a potential factor contributing to inter-doctor variability in the use of laboratory tests.

Given a detailed breakdown of the age of patients on the individual practice lists, we examined the relationship between the percentage of patients over age 65 and ordering rates for electrolyte, thyroid, liver, and lipid profiles. In addition, we investigated the link between gender and requesting activity for PSA (for males) and thyroid profiles (for females) and evaluated test requesting patterns for diabetes mellitus and their relationship to QOF clinical indicator scores in diabetes.

### Outcome measures

Our outcome of interest was to identify the presence and extent of variations in primary care laboratory tests ordering and to evaluate temporal changes in both the standardised number of test requests and between-practice variability in requesting. In addition, we studied demographic, socio-economic, geographical, and clinical factors that may explain this variation.

### Statistical analysis

Inter-practice variability in test requests was assessed by calculating the variance (σ^2^). Furthermore, we computed ‘variability index’ (Var_i_) defined as the top decile divided by the bottom decile of standardized test request rates.^10^ Var_i_ is a dimensionless measure of dispersion allowing us to compare the amount of variation in request rates of individual tests despite their differences in scale. The Shapiro-Wilk test of normality was used to determine if the distribution of test ordering data deviated from a normal distribution.^22^ Since the distribution of laboratory test request rates was found to be non-normal, the non-parametric statistics were implemented to perform further analysis. Mann Whitney U (MWU) test was employed to compare distributions of laboratory test rates between GP practices located in rural and urban areas.^23^ The homogeneity of variances for requesting activity in rural and urban GP practices was assessed with the Fligner-Killeen (FK) test.^24^ Significance of temporal changes in median and variability in test requesting rates was examined with the Mann–Kendall (MK) test.^25^ In all conducted tests, a *p* < 0.05 was considered significant. Kendall rank correlation coefficient (τ)^26^ was calculated to test the relationship between adjusted requesting rates and 1) Multiple Deprivation Measure; 2) Health Deprivation & Disability Deprivation rank; 3) proportions of patients over age 65; 4) distribution of patient’s gender; and 5) QOF clinical indicator scores. The Kendall coefficient (−1 ≤ τ ≤ 1) measures the strength of a correlation between two variables and assesses the degree of overall correspondence of variables’ ranking.^26^

### Patient and public involvement

No patients or general practitioners were involved in defining the research question or outcome measures nor were they engaged in the design or implementation this study. The study was approved by the Western Health and Social Care Trust (WHSCT). All identifiable information used for the purpose of the study was anonymised and not traceable to individual patients or general practitioners.

## Results

### Test ordering patterns

We analysed the laboratory test request rates of 55 general practices within the catchment area of the WHSCT comprising a total of 316 382 (2011-12), 316 688 (2012-13), 318 057 (2013-14), 319 383 (2014-2015), and 326 429 (2015-16) patients. Figure 1 shows the total number of 8 considered clinical biochemistry tests/test groups ordered during the study period of five consecutive years. The total number of ordered tests was 523 111 (2011-12), 531 849 (2012-13), 531 583 (2013-14), 525 146 (2014-2015), and 542 118 (2015-16) with electrolyte profiles found to be the most frequently ordered tests, making up approximately 30% of all requests in each year.

**Figure 1.**
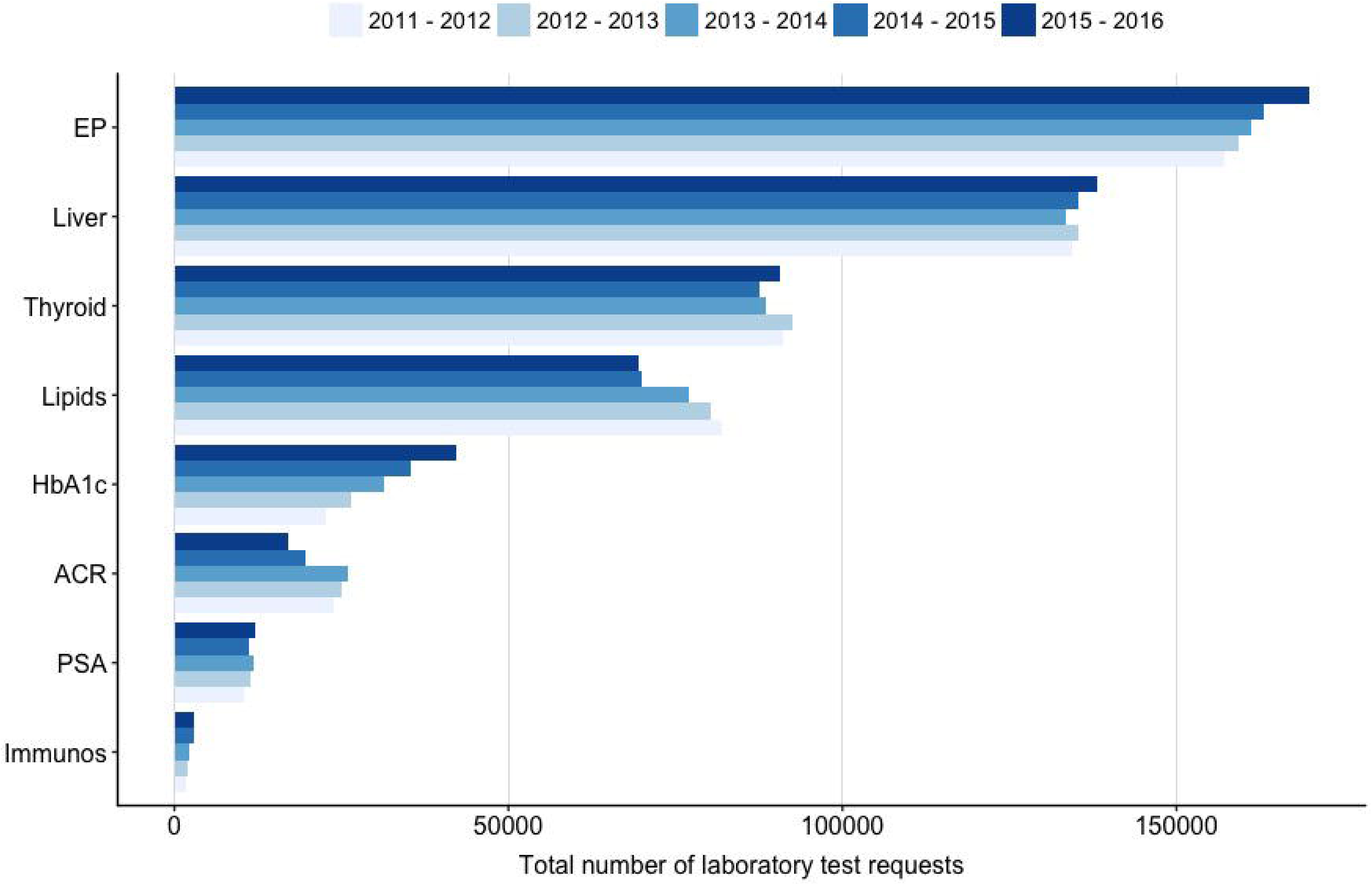
The total number of 8 considered clinical biochemistry tests/test groups ordered by 55 general practices during the period 1 Apr 2011 – 31 Mar 2016.

We observed substantial between-practice variability for all studied tests (table 1). In the considered period, the Var_i_ was lowest for electrolyte profiles (2.0-2.3), liver profiles (2.2-2.5), and HbA_1c_ (2.2-2.9) and highest for immunoglobulins reaching the value of 69.8 in 2012-13. The large variability was caused by several GP practices with abnormally high/low ordering rates of laboratory tests (figure 2). For the majority of tests, the median of test request rates was found to be generally lower than the mean, suggesting that the distribution of ordered test was primarily affected by the presence of GP practices with extremely high numbers of standardized requests.

**Table 1.**
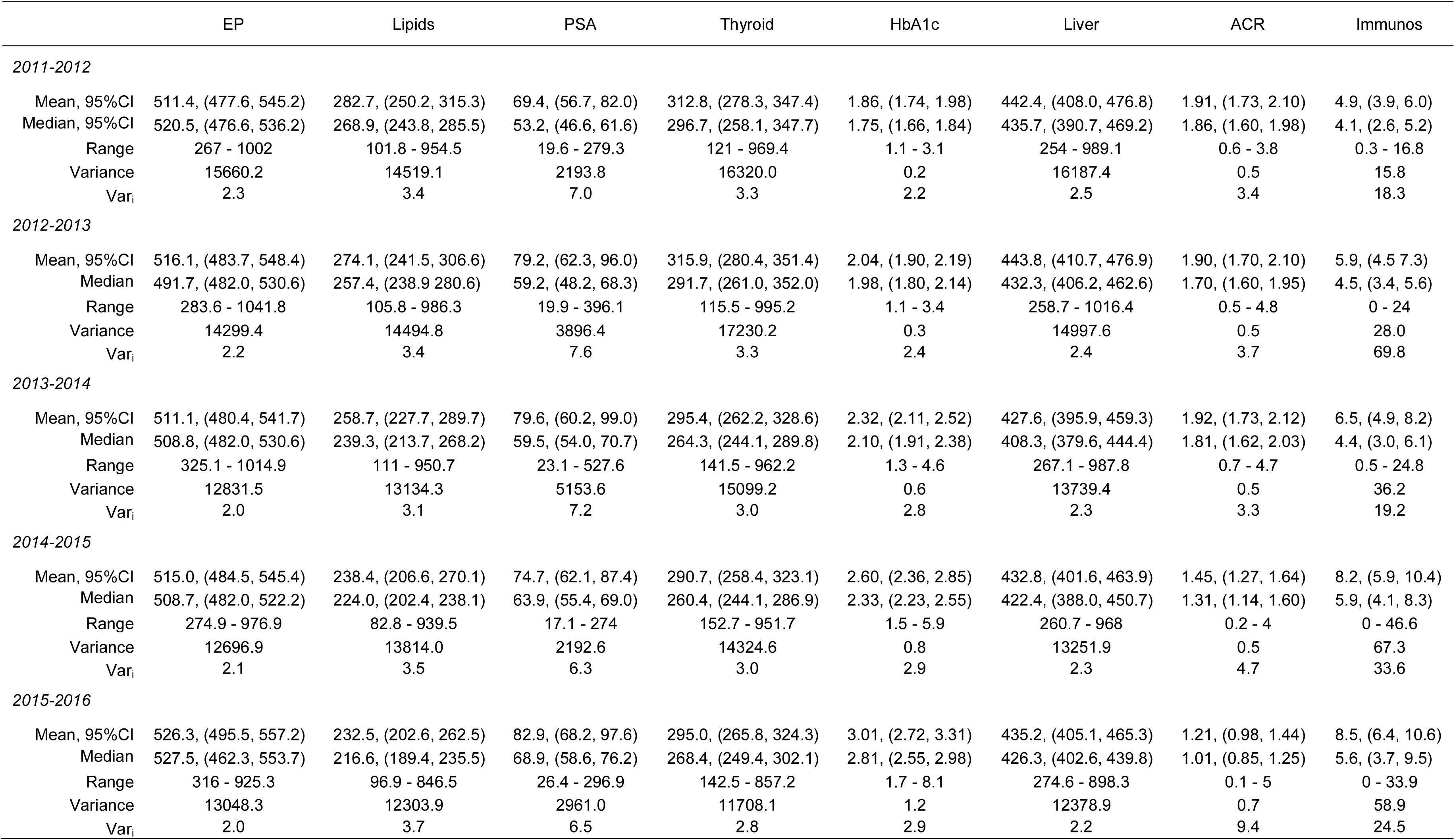
Overview of general statistics

|  | EP | Lipids | PSA | Thyroid | HbA1c | Liver | ACR | Immunos |
| --- | --- | --- | --- | --- | --- | --- | --- | --- |
| *2011-2012* |  |  |  |  |  |  |  |  |
| Mean, 95%CI | 511.4, (477.6, 545.2) | 282.7, (250.2, 315.3) | 69.4, (56.7, 82.0) | 312.8, (278.3, 347.4) | 1.86, (1.74, 1.98) | 442.4, (408.0, 476.8) | 1.91, (1.73, 2.10) | 4.9, (3.9, 6.0) |
| Median, 95%CI | 520.5, (476.6, 536.2) | 268.9, (243.8, 285.5) | 53.2, (46.6, 61.6) | 296.7, (258.1, 347.7) | 1.75, (1.66, 1.84) | 435.7, (390.7, 469.2) | 1.86, (1.60, 1.98) | 4.1, (2.6, 5.2) |
| Range | 267 - 1002 | 101.8 - 954.5 | 19.6 - 279.3 | 121 - 969.4 | 1.1 - 3.1 | 254 - 989.1 | 0.6 - 3.8 | 0.3 - 16.8 |
| Variance | 15660.2 | 14519.1 | 2193.8 | 16320.0 | 0.2 | 16187.4 | 0.5 | 15.8 |
| Var<sub>i</sub> | 2.3 | 3.4 | 7.0 | 3.3 | 2.2 | 2.5 | 3.4 | 18.3 |
| *2012-2013* |  |  |  |  |  |  |  |  |
| Mean, 95%CI | 516.1, (483.7, 548.4) | 274.1, (241.5, 306.6) | 79.2, (62.3, 96.0) | 315.9, (280.4, 351.4) | 2.04, (1.90, 2.19) | 443.8, (410.7, 476.9) | 1.90, (1.70, 2.10) | 5.9, (4.5, 7.3) |
| Median | 491.7, (482.0, 530.6) | 257.4, (238.9, 280.6) | 59.2, (48.2, 68.3) | 291.7, (261.0, 352.0) | 1.98, (1.80, 2.14) | 432.3, (406.2, 462.6) | 1.70, (1.60, 1.95) | 4.5, (3.4, 5.6) |
| Range | 283.6 - 1041.8 | 105.8 - 986.3 | 19.9 - 396.1 | 115.5 - 995.2 | 1.1 - 3.4 | 258.7 - 1016.4 | 0.5 - 4.8 | 0 - 24 |
| Variance | 14299.4 | 14494.8 | 3896.4 | 17230.2 | 0.3 | 14997.6 | 0.5 | 28.0 |
| Var<sub>i</sub> | 2.2 | 3.4 | 7.6 | 3.3 | 2.4 | 2.4 | 3.7 | 69.8 |
| *2013-2014* |  |  |  |  |  |  |  |  |
| Mean, 95%CI | 511.1, (480.4, 541.7) | 258.7, (227.7, 289.7) | 79.6, (60.2, 99.0) | 295.4, (262.2, 328.6) | 2.32, (2.11, 2.52) | 427.6, (395.9, 459.3) | 1.92, (1.73, 2.12) | 6.5, (4.9, 8.2) |
| Median | 508.8, (482.0, 530.6) | 239.3, (213.7, 268.2) | 59.5, (54.0, 70.7) | 264.3, (244.1, 289.8) | 2.10, (1.91, 2.38) | 408.3, (379.6, 444.4) | 1.81, (1.62, 2.03) | 4.4, (3.0, 6.1) |
| Range | 325.1 - 1014.9 | 111 - 950.7 | 23.1 - 527.6 | 141.5 - 962.2 | 1.3 - 4.6 | 267.1 - 987.8 | 0.7 - 4.7 | 0.5 - 24.8 |
| Variance | 12831.5 | 13134.3 | 5153.6 | 15099.2 | 0.6 | 13739.4 | 0.5 | 36.2 |
| Var<sub>i</sub> | 2.0 | 3.1 | 7.2 | 3.0 | 2.8 | 2.3 | 3.3 | 19.2 |
| *2014-2015* |  |  |  |  |  |  |  |  |
| Mean, 95%CI | 515.0, (484.5, 545.4) | 238.4, (206.6, 270.1) | 74.7, (62.1, 87.4) | 290.7, (258.4, 323.1) | 2.60, (2.36, 2.85) | 432.8, (401.6, 463.9) | 1.45, (1.27, 1.64) | 8.2, (5.9, 10.4) |
| Median | 508.7, (482.0, 522.2) | 224.0, (202.4, 238.1) | 63.9, (55.4, 69.0) | 260.4, (244.1, 286.9) | 2.33, (2.23, 2.55) | 422.4, (388.0, 450.7) | 1.31, (1.14, 1.60) | 5.9, (4.1, 8.3) |
| Range | 274.9 - 976.9 | 82.8 - 939.5 | 17.1 - 274 | 152.7 - 951.7 | 1.5 - 5.9 | 260.7 - 968 | 0.2 - 4 | 0 - 46.6 |
| Variance | 12696.9 | 13814.0 | 2192.6 | 14324.6 | 0.8 | 13251.9 | 0.5 | 67.3 |
| Var<sub>i</sub> | 2.1 | 3.5 | 6.3 | 3.0 | 2.9 | 2.3 | 4.7 | 33.6 |
| *2015-2016* |  |  |  |  |  |  |  |  |
| Mean, 95%CI | 526.3, (495.5, 557.2) | 232.5, (202.6, 262.5) | 82.9, (68.2, 97.6) | 295.0, (265.8, 324.3) | 3.01, (2.72, 3.31) | 435.2, (405.1, 465.3) | 1.21, (0.98, 1.44) | 8.5, (6.4, 10.6) |
| Median | 527.5, (462.3, 553.7) | 216.6, (189.4, 235.5) | 68.9, (58.6, 76.2) | 268.4, (249.4, 302.1) | 2.81, (2.55, 2.98) | 426.3, (402.6, 439.8) | 1.01, (0.85, 1.25) | 5.6, (3.7, 9.5) |
| Range | 316 - 925.3 | 96.9 - 846.5 | 26.4 - 296.9 | 142.5 - 857.2 | 1.7 - 8.1 | 274.6 - 898.3 | 0.1 - 5 | 0 - 33.9 |
| Variance | 13048.3 | 12303.9 | 2961.0 | 11708.1 | 1.2 | 12378.9 | 0.7 | 58.9 |
| Var<sub>i</sub> | 2.0 | 3.7 | 6.5 | 2.8 | 2.9 | 2.2 | 9.4 | 24.5 |

**Figure 2.**
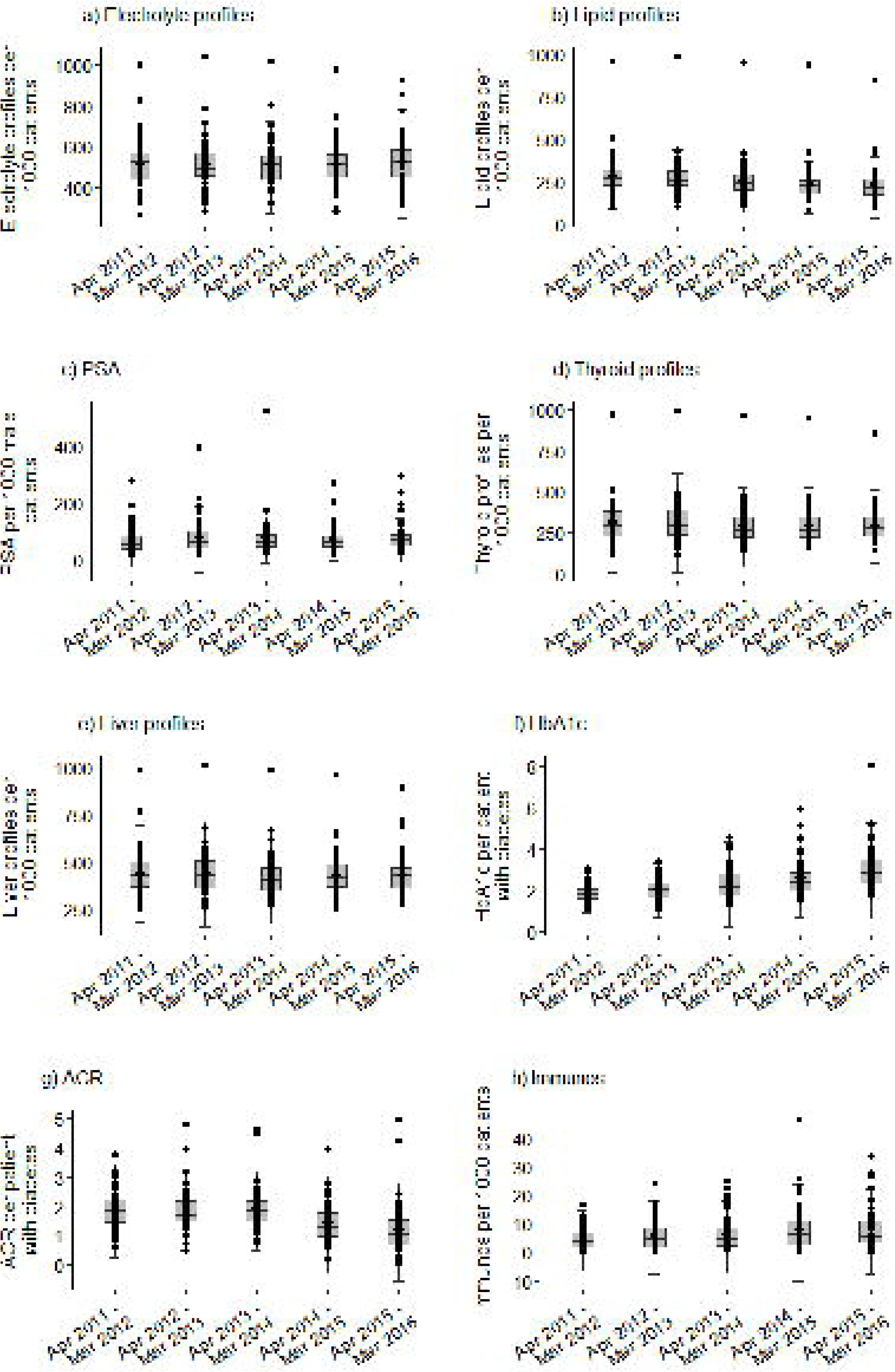
Temporal variability of the standardized laboratory test requests for (a) electrolyte profiles, (b) lipid profile, (c) prostate-specific antigen (PSA), (d) thyroid profiles, (e) liver profiles, (f) HbA_1c_, (g) urine albumin/creatinine ratio (ACR), and (h) immunoglobulins (Immunos) for 55 considered general practices. Each data point (dot): a single practice. Solid, horizontal line inside the box: median. Lower and upper “hinges” of the boxplots: 1^st^ and 3^rd^ quartiles, respectively. Lower and upper extremes of whiskers: interval boundaries of the non-outliers (black dots). Data outside interval: outliers.

### Changes in the number and variation of test requests over time

Over the period of investigation, we observed an increase in the median standardized number of electrolyte profiles, immunoglobulins, PSA, and HbA_1c_ (figure 3); however, the Mann–Kendall (MK) test indicated the statistically significant upward trend only for requesting rates of PSA (*p*_*MK*_ = 0.03) and HbA_1c_ (*p*_*MK*_ = 0.03). For lipid profiles, thyroid profiles, liver profiles, and ACR, we reported the overall decline of the median number of test request rates. Yet, the downward trend was found statistically significant only for lipid profiles (*p*_*MK*_ = 0.03).

**Figure 3.**
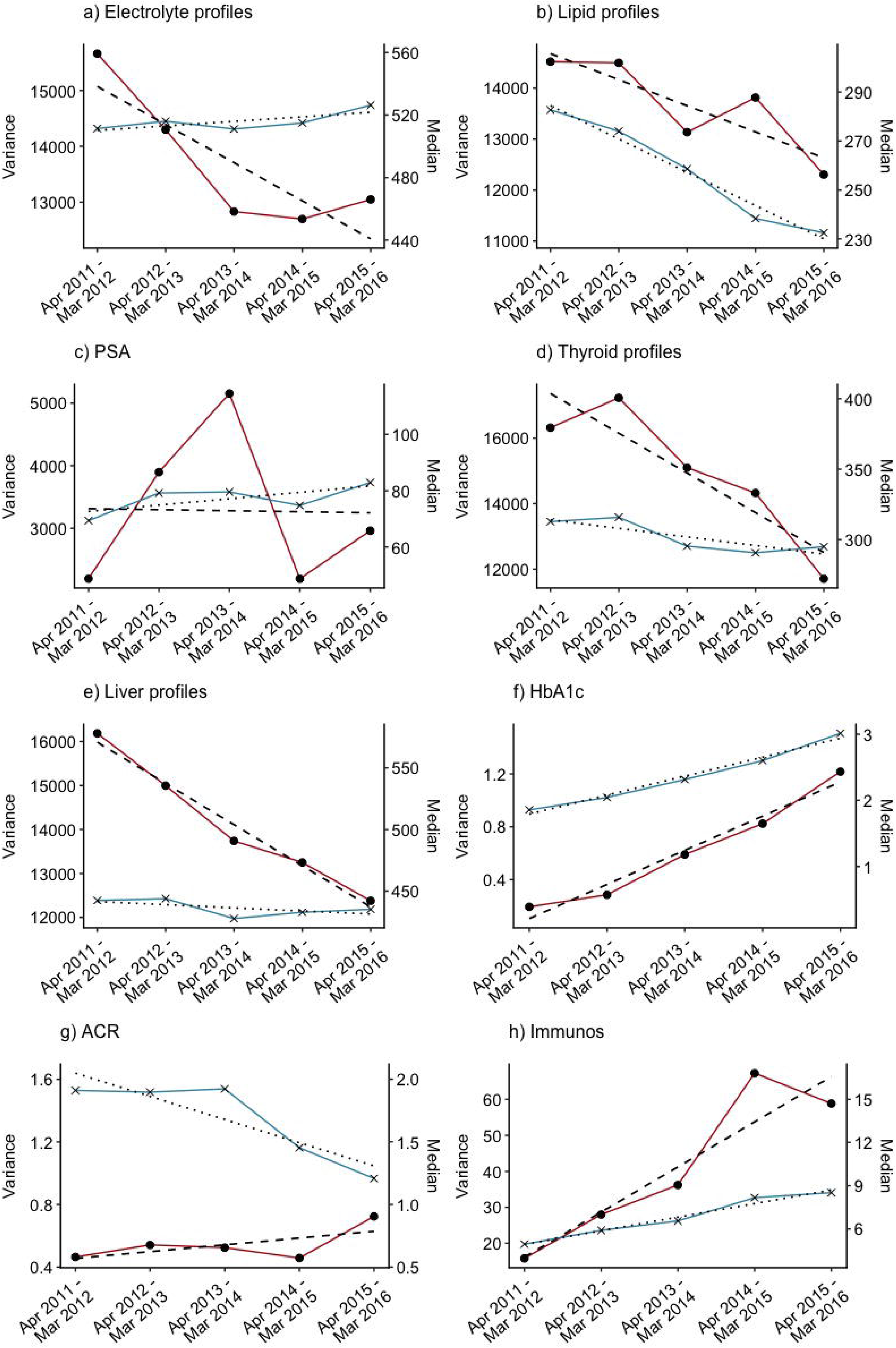
Trend lines for median (blue) and variance (red) of the standardized number of (a) electrolyte profiles, (b) lipid profile, (c) prostate-specific antigen (PSA), (d) thyroid profiles, (e) liver profiles, (f) HbA_1c_, (g) urine albumin/creatinine ratio (ACR), and (h) immunoglobulins (Immunos).

The variance for electrolyte profiles, lipid profiles, thyroid profiles, and liver profiles requests fell by 16.8%, 15.3%, 28.3%, and 23.5% respectively (figure 3). The increase in variance was observed for immunoglobulins (272%), ACR (40%), and HbA_1c_ (500%). Nonetheless, only the increase in variance of HbA_1c_ requests (*p*_*MK*_ = 0.03) and decrease in variance of standardized number of liver profiles (*p*_*MK*_ = 0.03) were found significant at the 95% confidence level. For PSA test request rates, we observed fluctuations in variance with no clear trend.

### Test ordering variation between general practices and explanatory factors

The relationship between test requesting rates and potential explanatory factors was established based on the information on the number of ordered tests, patient demographics, and Quality and Outcomes Framework indicator scores obtained for the period 1 Apr 2015 – 31 Mar 2016.

#### Demographic characteristics of patient population

Proportion of the oldest age category of patients (> 75) constituted a relatively small group in each GP practice (mean = 6.1%, 95%CI = (5.7%,6.5%)). Hence, we combined the 65-74 and > 75 age categories to create a more meaningful group of patients in the older age band. We examined four tests for which differences in proportions of older patients were expected to be reflected in test requesting i.e. electrolyte profiles, liver profiles, lipid profiles, and thyroid profiles. Since we were unable to obtain a consolidated data on both gender and age of patients, we did not assess the relationship between the PSA requesting rates and the category of males aged 65 and over. We found a very weak correlation between the adjusted request rates of selected laboratory tests and the percentage distribution of patients of age over 65, with τ ranging from 0.08 for thyroid profiles to 0.23 for lipid profiles (supplementary table S1). Since previous studies reported on a higher rate of testing of thyroid hormones in females,^27^ we looked at the strength of a relationship between the percentage of females in individual GP practices and requesting rates for thyroid profiles. We found a weak, in fact negative, association between these two characteristics, with τ = −0.2 (supplementary figure S2).

In addition, we examined the effect of gender distribution on the standardized number of PSA tests, with the PSA ordering rates expected to be higher for GP practices with higher percentage of males (supplementary figure S2). The coefficient τ = 0.2 implied a weak degree of correlation between these two variables. Note that we acknowledge the fact that a combined effect of sex and age distributions might have had a more significant effect on PSA requesting activity. However, we were unable to extract such data.

#### Practice setting

Figure 4 shows a temporal median-variance relationship of the standardized number of laboratory test requests for rural and urban areas. Large differences were identified for PSA, lipid profiles, thyroid profiles, and liver profiles. The median number of tests ordered annually by practices located in rural areas was higher by approximately 27-37% for PSA, 14-30% for lipid profiles, 14-38% for thyroid profiles, and 8-23% for liver profiles. For ACR and immunoglobulins the median requesting rates were lower in rural areas by 1-27% and 18-57% respectively. Across five consecutive time periods, Mann Whitney U (MWU) test showed significant differences between ‘rural’ and ‘urban’ distributions of laboratory test rates for lipid profiles (*p*_*MWU*_ ranging from 0.00004 to 0.03) and PSA (*p*_*MWU*_ between 0.003 and 0.03). The significant rural-urban differences in requesting activity of thyroid and liver profies (*p*_*MWU*_ < 0.05) were observed only in the first two years of the study (table 2).

**Table 2.** The significance of differences in the distribution and variability in test request rates between GP practices located in rural and urban areas. *p*_*MWU*_: Mann Whitney U *p*-value assessing differences in distributions of laboratory test rates between rural and urban practices. *p*_*FK*_: Fligner-Killeen *p*-value referring to the significance level of differences in variances. For both tests, a *p* < 0.05 was considered significant (*).

|  | Electrolyte<br>profiles | Lipid<br>profiles | PSA | Thyroid<br>profiles | HbA1c | Liver<br>profiles | ACR | Immunoglobulins |
| --- | --- | --- | --- | --- | --- | --- | --- | --- |
| $p_{MWU}$ | | | | | | | | |
| 2011-2012 | 0.1 | 0.0004* | 0.01* | 0.009* | 0.3 | 0.03* | 0.4 | 0.2 |
| 2012-2013 | 0.08 | 0.001* | 0.03* | 0.01* | 0.7 | 0.02* | 0.5 | 0.4 |
| 2013-2014 | 0.8 | 0.01* | 0.02* | 0.07 | 1.0 | 0.3 | 0.5 | 0.4 |
| 2014-2015 | 0.6 | 0.01* | 0.02* | 0.1 | 0.9 | 0.3 | 0.5 | 0.03* |
| 2015-2016 | 0.7 | 0.03* | 0.003* | 0.07 | 0.7 | 0.3 | 0.08 | 0.01* |
| $p_{FK}$ | | | | | | | | |
| 2011-2012 | 0.4 | 0.5 | 0.03* | 0.2 | 0.6 | 1 | 0.2 | 0.8 |
| 2012-2013 | 0.3 | 0.6 | 0.01* | 0.3 | 0.9 | 0.4 | 0.7 | 0.3 |
| 2013-2014 | 0.2 | 0.6 | 0.06 | 0.03* | 0.5 | 0.7 | 1 | 0.5 |
| 2014-2015 | 0.9 | 0.6 | 0.1 | 0.02* | 0.5 | 0.9 | 0.3 | 0.6 |
| 2015-2016 | 0.7 | 0.7 | 0.2 | 0.2 | 0.2 | 0.7 | 0.8 | 0.1 |

**Figure 4.**
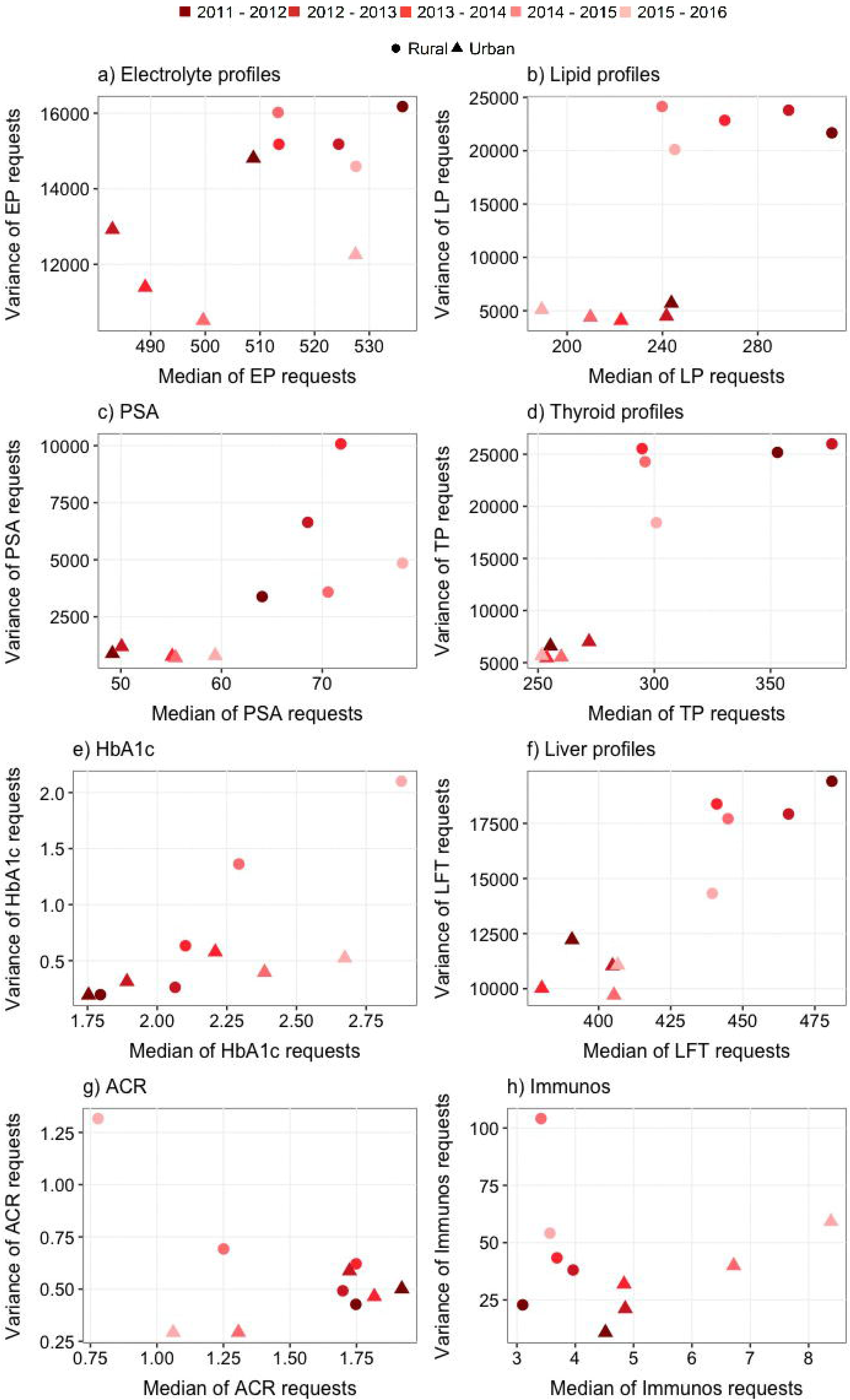
The temporal median-variance relationship of the standardized number of laboratory test requests across the years. Circle/triangle: rural/urban general practices.

Rural practices had significantly higher variability in test requesting than urban practices at all five time points. The variance in requesting rates in rural GP practices was higher by 9-52% for electrolyte profiles, 280-460% for lipid profiles, 280-1212% for PSA, 224-367% for thyroid profiles, 29-82% for liver profiles, 2-301% for HbA_1c_ (except for the period 2012-13 when the variance was 16% lower in rural areas), 34-50% for ACR (except for the periods of 2011-12 and 2012-13 when the variance in rural areas was lower by 15% and 16% respectively), and 36-161% for immunoglobulins (except for the period of 2015-16 when the variance was 9% lower in rural areas). Furthermore, we found the rural-urban differences in variance for ordering rates of PSA and thyroid profiles statistically significant (*p*_*FK*_ < 0.05) (table 2)

#### Socioeconomic factors

We observed a very weak association between the Multiple Deprivation Measure (MDM) and requesting activity. Accordingly, the lowest Kendall τ of 0.01 was reported for HbA_1c_ while the highest τ = 0.18 for PSA (supplementary table S3). Similarly, the relationship between the Health Deprivation & Disability rank and the standardized number of test request was the lowest for HbA_1c_ (τ = 0.06) and highest for PSA (τ = 0.31). Again, the τ value indicated a weak correlation in both cases.

#### Quality and Outcomes Framework indicators

To evaluate the relationship between standardized number of laboratory tests and QOF indicator scores, we looked at the management of diabetes. There are two main reasons for that. First, the guidelines for management of diabetes are widely used by practitioners to guide the care of their patients. Secondly, HbA_1c_ is a specific test in diabetes care and does not play an important role in the monitoring or diagnosis of any other condition.

The three target levels for HbA_1c_ (59, 64, and 75 mmol/mol) in the QOF were introduced to improve glycaemic control across the distribution of values. We investigated the overall QOF points under these 3 categories i.e. DM007 (‘The percentage of patients with diabetes in whom the last HbA_1c_ is 59 mmol/mol or less in the preceding 15 months’), DM008 (‘The percentage of patients with diabetes in whom the last HbA_1c_ is 64 mmol/mol or less in the preceding 15 months’), and DM009 (‘The percentage of patients with diabetes in whom the last HbA_1c_ is 75 mmol/mol or less in the preceding 15 months’). We also combined the number of points for each of these 3 categories, calculated the overall achievement rate, and investigated its relationship with inter-practitioner variability in test requesting of HbA_1c_.

All practices achieved the maximum 17 points available under QOF clinical indicator DM007. All but nine practices attained the maximum 8 points available under DM008 (range: 4.1-8.0) and 31 practices attained the maximum 10 points available under DM009 (range: 6.36-10.0). The strength of relationship between the number of HbA_1c_ tests performed and the GP practice effectiveness, as measured by the QOF overall achievement rate (combined DM007, DM008, and DM009) was very weak (τ = 0.12).

## Discussion

We evaluated temporal changes in variability and number of laboratory tests ordered by individual GP practices in the WHSCT. We also investigated a range of key demographic, socioeconomic, geographical, and clinical factors to assess whether any of these factors are likely to explain a part of the observed variation in requesting activity.

While we found an overall decrease in the median request rates for lipid profiles, thyroid profiles, liver profiles, and ACR (statistically significant only for lipid profiles), we observed an increase in the median standardized number of PSA, HbA_1c_, immunoglobulins, and electrolyte profiles (statistically significant only for PSA and HbA_1c_). Furthermore, we identified substantial differences in variation of adjusted test request rates for individual tests, with an index of variability oscillating between 2.0-2.9 for electrolyte profiles, liver profiles, and HbA_1c_, 2.8-3.3 for thyroid profiles, 3.1-3.7 for lipid profiles, 6.3-7.6 for PSA, and reaching up to 69.8 for immunoglobulins. Our finding of high levels of variability between practices is consistent with previous research.^6,10,28,29^ This large variation may suggest that a portion of ordered tests had been not clinically relevant according to standards and guidelines and therefore, had limited or no benefit to the patient care.

Temporal changes in between practice variability of request rates were associated with a reduction in variance for electrolyte profiles (16.8%), lipid profiles (15.3%), thyroid profiles (28.3%), and liver profiles (23.5%) and the increase in variation of ordering rates for ACR (40%), immunoglobulins (272%), and HbA_1c_ (500%) over the period of investigation. The overall downward trend in variability of profile test request rates might have been related to the implementation of the demand optimization intervention in the WHSCT over the three year period from Apr 2012 to Mar 2015. The intervention was developed to support appropriate use of laboratory testing in the WHSCT and consisted of several elements including educational materials on the benefits of the optimal tests utilization, information on minimal retest intervals, the review of the test requesting processes, and financial incentive. The 500% increase in variability of HbA_1c_ ordering rates, on the other hand, may have been caused by between practice differences in the adoption of HbA_1c_ as a diagnostic test (from 2012) and inconsistent implementation of new guidelines on appropriate rate of diabetes monitoring.

The inter-practitioner variations in test ordering were found unrelated to demographic and socioeconomic factors. We showed that distributions of patients’ age and sex had little impact on ordering rates for a predetermined set of pathology tests. The socioeconomic status of GP practice also did not appear to identify low or high requestors of laboratory tests. No clear relationship between test ordering and age, gender or deprivation measures was reported by other studies.^6,30-32^

The practice location was found to be a significant factor associated with variability in test use. Both the variability and median number of request rates were generally higher in GP practices located in rural areas at all time points. This probably reflects the differences in clinical practice associated with the specific aspects of practice organisation and workflow in rural and urban practices as well as personal characteristics of general practitioners. The practice location was previously related to the between-practice variation in prescribing.^33-35^ Further studies are needed to explore in more detail the potential reasons for the rural-urban discrepancy in test utilization.

Finally, we found no evidence of a significant correlation between test requesting rates and clinical outcomes. This observation is consistent with other studies.^10^ The fact that the majority of practices attained maximum points for the three target levels for HbA_1c_ (59, 64, and 75 mmol/mol) implies that the glycaemic control was generally good and could not explain the variability in HbA_1c_ testing.

### Implications for practice and research

The findings of our study suggest that there may be considerable potential for the rationalization of test ordering through minimizing the between practice variability in test utilization. Yet, it is important to establish whether the observed variation is, and to what extent, associated with over-requesting (unnecessary repeat requesting of tests) in GP practices with high ordering rates or on the contrary, it reflects a failure to prescribe clinically indicated tests by GP practices characterized by low use of tests. Further analysis on the degree and detection of inappropriate use of laboratory resources in primary care could contribute to improving the consistency, efficiency, and cost-effectiveness of patient diagnostics, monitoring, and treatment and hence, reduce unnecessary costs to patient care.

Further exploration of variations in requesting activity in the context of factors not considered in this study (e.g. practitioner-specific characteristics) may help identify and implement appropriate optimization strategies to manage demand for laboratory tests. Previous studies reported on the number of successful approaches in fostering best practice and reducing variation in test utilization including implementing locally agreed clinical guidelines, changing test order forms, incorporating clinical decision support tools with embedded retest interval rules, conducting audits of GPs on their request rates, and providing financial incentives.^36-42^ In fact, several studies showed that multifaceted interventions were most successful in optimizing laboratory demand.^43,44^

Finally, it is essential to better understand implications of variability in laboratory test requesting for the cost and quality of care. In particular, an important question to be answered is whether or not the growing costs associated with increased use of laboratory services has led to commensurate benefits to the patient.

### Strengths and limitations

Test requesting data were directly extracted from the HSC Business Object system that captures information on tests’ use from three clinical chemistry departments of Altnagelvin Area Hospital, Tyrone County Hospital, and South West Acute Hospital. Accordingly, our analysis was based on all available data regarding requesting activity in the N. Ireland Western Health and Social Care Trust (WHSCT) in the period of 1 Apr 2011 – 31 Mar 2016 and hence, was not subject to selection bias. Furthermore, our study provides a first comprehensive insight into the use of laboratory tests and factors accounting for the variation in between practice test utilization in the WHSCT primary care system.

Due to data unavailability, we were unable to investigate the relationship between laboratory use patterns and practitioner-specific characteristics including GP’s age, education, and medical training. Such analysis could help identify potential reasons behind variation in clinical practice. Besides the lack of information on the practitioners, the present study was limited by a paucity of research evidence in this area. We were also unable to retrieve consolidated data on both gender and age of patients registered in individual GP practices and therefore, assess the combined effect of sex and age distributions on test requesting activity.

## Conclusion

This study investigated the patterns and temporal changes in request rates across a range of frequently ordered laboratory tests. In addition, it explored potential factors of the substantial inter-practice variability in the use of laboratory tests and found that differences in requesting activity appear unrelated to either demographic and socioeconomic characteristics of GP practices or clinical outcome indicators. Our results highlight the need for further investigations to identify other potential factors that may account for the differences in test utilization between practitioners.

## Acknowledgments

The authors would wish to thank Dr KongFatt Wong-Lin for helpful discussions and Graham Moore for computing and technical support.

## Contributors

MB and MJO had the original idea for this study. SA led the data collection. MB designed the methodology, performed the analysis, and drafted the manuscript. MJO, CM, and LM contributed to the drafting and critical revision of the manuscript.

## Ethics approval

Not applicable.

## Data sharing statement

No additional data are available.

